# Widespread decrease of cerebral vimentin-immunoreactive astrocytes in depressed suicides

**DOI:** 10.1101/2020.12.12.422510

**Authors:** Liam Anuj O’Leary, Claudia Belliveau, Maria Antonietta Davoli, Jie Christopher Ma, Arnaud Tanti, Gustavo Turecki, Naguib Mechawar

**Affiliations:** McGill Group for Suicide Studies, Douglas Mental Health University Institute, Verdun, QC, Canada; Integrated Program in Neuroscience, McGill University, Montreal, Canada; Department of Psychiatry, McGill University, Montreal, QC, Canada

**Keywords:** human, post-mortem, depression, suicide, astrocyte, vimentin, GFAP

## Abstract

Postmortem investigations have implicated cerebral astrocytes immunoreactive (-IR) for glial fibrillary acidic protein (GFAP) in the etiopathology of depression and suicide. However, it remains unclear whether astrocytic subpopulations IR for other astrocytic markers are similarly affected. Astrocytes IR to vimentin (VIM) display different regional densities than GFAP-IR astrocytes in the healthy brain, and so may be differently altered in depression and suicide. To investigate this, we compared the densities of GFAP-IR astrocytes and VIM-IR astrocytes in postmortem brain samples from depressed suicides and matched non-psychiatric controls in three brain regions (dorsomedial prefrontal cortex, dorsal caudate nucleus and mediodorsal thalamus). A quantitative comparison of the fine morphology of VIM-IR astrocytes was also performed in the same regions and subjects. Finally, given the close association between astrocytes and blood vessels, we also assessed densities of CD31-IR blood vessels. Like for GFAP-IR astrocytes, VIM-IR astrocyte densities were found to be globally reduced in depressed suicide relative to controls. By contrast, CD31-IR blood vessel density and VIM-IR astrocyte morphometric features in these regions were similar between groups, except in prefrontal white matter, in which vascularization was increased and astrocytes displayed fewer primary processes. By revealing a widespread reduction of cerebral VIM-IR astrocytes in cases vs. controls, these findings further implicate astrocytic dysfunctions in depression and suicide.

## 2 Introduction

Astrocytes were first identified as a glial cell type in the human brain more than a hundred years ago, and until a few decades ago were mostly seen to have a passive role of providing nutritional support for neurons (Andriezen, 1893). Animal studies have since revealed that astrocytes can strongly modulate most facets of neuronal activity, including neuronal firing, neurotransmitter synthesis, neurotransmitter reuptake, and synaptic transmission (Norenberg & Martinez-Hernandez, 1979; Rothstein *et al*., 1996; Duan *et al*., 1999; Cui *et al*., 2018; Covelo & Araque, 2018). Astrocytes might especially influence neuronal activity in the human brain, as they are almost threefold larger in volume and fourfold faster at signaling in the human cortex than in the mouse cortex (Oberheim *et al*., 2009; O’Leary *et al*., 2020).

The first postmortem investigations of major depressive disorder (MDD) reported reduced glial (but not neuronal) densities in the ventral anterior cingulate cortex (Öngür et al., 1998), the orbitofrontal cortex (Rajkowska et al., 1999), and the amygdala (Bowley et al., 2002). These findings were later attributed to a reduced number of cerebral astrocytes, particularly those immunoreactive (-IR) for the astrocyte-specific marker glial fibrillary acidic protein (GFAP). Postmortem brain samples from depressed individuals have fewer GFAP-IR astrocytes (Miguel-Hidalgo et al., 2000; Webster et al., 2001; Si et al., 2004; Altshuler et al., 2010; Cobb et al., 2016; Rajkowska et al., 2018), and lower levels of GFAP mRNA and protein (Chandley et al., 2013; Fatemi et al., 2004; Torres-Platas et al., 2016). Of all psychiatric conditions, GFAP is most implicated in MDD (Kim et al., 2018). However, GFAP labels only a minority of astrocytes, and so may misrepresent astrocytic phenotypes in MDD (Sofroniew & Vinters, 2010). For instance, the hippocampal CA1 region has lower densities of S100B-IR, but not GFAP-IR, astrocytes in MDD (Gos et al., 2013). Conversely, the amygdala has lower densities of GFAP-IR, but not S100B-IR, astrocytes in MDD (Altshuler et al., 2010; Hamidi et al., 2004). Hence, GFAP expression might not capture the widespread phenotype of astrocyte dysfunction in postmortem studies of MDD, including morphometric differences in cortical fibrous astrocytes (Torres-Platas et al., 2011), reduced vascular coverage (Rajkowska et al., 2013), differential methylation patterns for genes enriched in astrocytes (Nagy et al., 2015), and abnormally low expression levels of astrocytic glutamate transporters (Chandley et al., 2013; Nagy et al., 2015; Zhao et al., 2016), and gap junction proteins (Ernst et al., 2011; Miguel-Hidalgo et al., 2014; Tanti et al., 2019).

Like GFAP, vimentin (VIM) is a type III intermediate filament that is strongly expressed in cerebral astrocytes, however, it is also expressed in vascular endothelial cells. The study of VIM-IR astrocytes has been relative rare, especially given the functional relationship between VIM and GFAP proteins. For instance, VIM has a reciprocal expression profile with GFAP during development, can functionally compensate for the transgenic loss of GFAP expression, and is peculiarly absent in Rosenthal fibres — a defining pathological feature of Alexander disease, a genetic condition associated with GFAP mutations (Dahl, 1981; Tomokane *et al*., 1991; Wilhelmsson *et al*., 2004). A previous postmortem study has found qualitative differences in VIM-IR astrocytes in neurological conditions (Yamada et al., 1992). In a recent postmortem study, we characterized VIM immunoreactivity in different cortical and subcortical brain regions using samples from healthy individuals having died suddenly, and found that VIM-IR astrocytes had different densities from GFAP-IR astrocytes, but that both GFAP-IR and VIM-IR astrocyte density inversely correlated with CD31-IR vascular density (O’Leary *et al*., 2020). We were then interested in how these findings might relate to depression.

Here, we compare the densities of GFAP-IR astrocytes, VIM-IR astrocytes and CD31-IR blood vessels in three brain regions from depressed suicides and matched non-psychiatric controls. In addition, we assessed VIM-IR astrocyte morphometry in MDD in the same sections. In depressed suicides, we found a general and consistent reduction in the density of GFAP-IR and VIM-IR astrocytes, as well as a significant increase in CD31-IR vascularization in the prefrontal cortex white matter. However, observed almost unaltered VIM-IR astrocyte morphometry in depressed suicides relative to controls. These findings indicate that in the brains of depressed individuals, regional variations in astrocyte densities are much stronger and widespread than changes in astrocyte morphometry.

## 3 Materials and Methods

### 3.1 Subjects and Tissue Processing

This study was approved by the Douglas Hospital Research Ethics Board. Brain samples were analyzed from adult Caucasian male depressed suicides (n=10) and non-psychiatric controls (n=10). Subject information is available in **Table 1**. All depressed suicides died during a major depressive episode. All controls died suddenly without any known inflammatory, psychiatric or neurological disorder. Brain donation and psychiatric diagnosis were as described previously (Torres-Platas et al., 2011). Subject groups were matched for three covariates: age (p = 0.75), tissue pH (p = 0.40), and postmortem interval (PMI; p = 0.38). For all 20 subjects, three brain regions were dissected from thick frozen sections: the dorsomedial prefrontal cortex (Brodmann Area (BA) 8/9), the dorsal caudate nucleus (precommissural) and the mediodorsal thalamus. We studied the PFC gray matter (PFC GM) and white matter (PFC WM) independently. Fresh-frozen 1cm^3^ tissue blocks from each region were fixed overnight in 10% formalin, suspended in 30% sucrose solution until equilibrium was reached, flash-frozen in −35°C isopentane, and cut on a sliding microtome into 50 µm-thick serial sections that were stored at −20°C in a cryoprotectant solution until processing for immunohistochemistry (IHC). Immunolabeling involved independently using each antibody listed in **Table 2** within a conventional DAB IHC protocol and a separate stereological series of sections, as described previously (O’Leary et al., 2020).

**Table 1:** Subject Information.

|  | Controls (n = 10) | Depressed Suicides (n = 10) |
| --- | --- | --- |
| | Mean ± SEM | Mean ± SEM |
| **Age** | 40.6 ± 5.0 | 38.5 ± 4.2 |
| **Sex** | 10M | 10M |
| **Tissue pH** | 6.5 ± 0.0 | 6.6 ± 0.0 |
| **PMI (h)** | 15.3 ± 2.8 | 21.8 ± 5.9 |
| **Cause of death** | 9 Natural, 1 Accidental | 10 Hanging |
| **Depression Status** | 0 | 10 |
| *Toxicology Report* |  |  |
| **Alcohol** | 1 | 3 |
| **Cocaine** | 0 | 2 |
| **Antidepressants** | 0 | 1 |

**Table 2:**
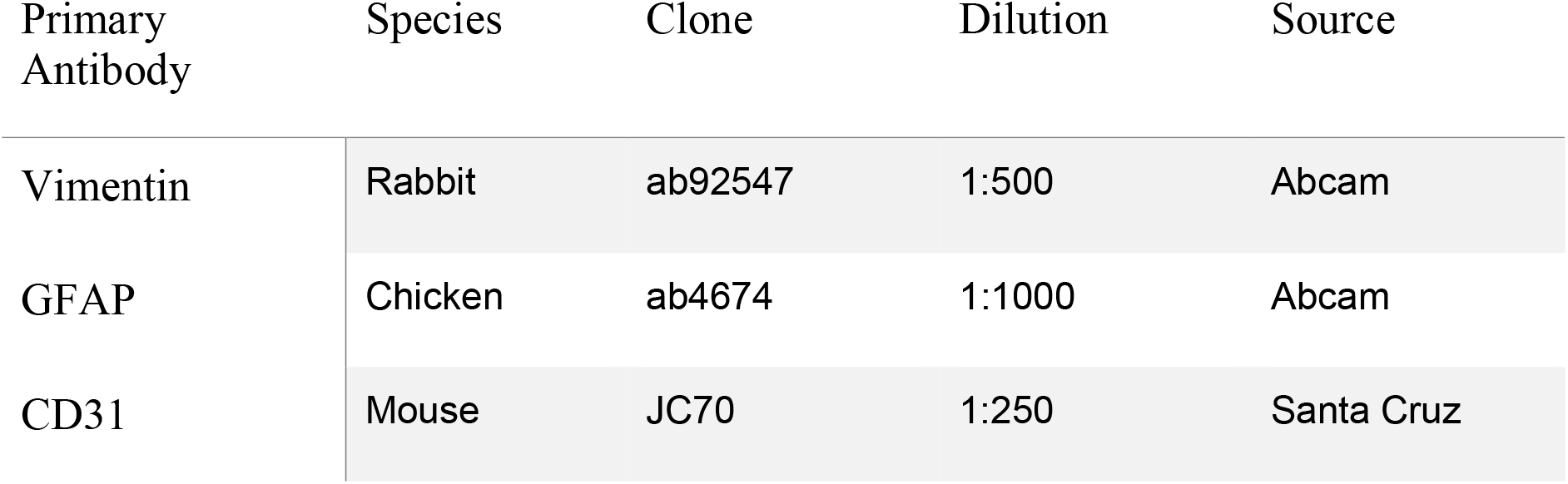
Antibodies: specifications.

| Primary Antibody | Species | Clone | Dilution | Source |
| --- | --- | --- | --- | --- |
| Vimentin | Rabbit | ab92547 | 1:500 | Abcam |
| GFAP | Chicken | ab4674 | 1:1000 | Abcam |
| CD31 | Mouse | JC70 | 1:250 | Santa Cruz |

### 3.2 Stereology and Morphology

Stereological cell counting and live tracing was performed as described previously (O’Leary et al., 2020), while additionally blinded to subject identities. Reliable stereological estimates, with a Gunderson coefficient of error (CE, m=1) < 0.10, were obtained by sampling the following percentage areas from section contours: 5% for GFAP in all regions; and 10% for VIM in the PFC GM and the caudate nucleus, 25% in the PFC WM, and 100% in the mediodorsal thalamus. A total of 320 VIM-IR astrocytes were reconstructed, four from the PFC GM, the PFC WM, the thalamus and the caudate nucleus of each subject.

### 3.3 Vascular Density

To estimate vascular density for its comparison with stereological cell densities, 15 brightfield images of CD31-IR vasculature were taken across four sections of a stereological series of sections for each region at low (10X objective) magnification. Using ImageJ software (NIH, USA), each image was converted into an 8-bit format and then manually thresholded to remove most background staining before the percentage area was measured. To reduce noise from background staining and artefacts, only ten values closest to the original mean value were used to calculate the mean % CD31-IR density for each region and subject.

### 3.4 Morphometric features

With the same slides and workstation used for stereological analysis, the morphometry of VIM-IR astrocytes were manually traced live using a 100X oil immersion objective and a computer-based tracing system (Neurolucida Explorer, MBF Bioscience). Immunostained cells were randomly selected, but had to display the following features in ordered to be selected for reconstruction: (1) unobstructed by neighboring cells; (2) of representative size and shape; (3) of equal staining across cellular compartments; (4) contained within the thickness of the section; (5) forming clear endfeet contacts with a VIM-IR blood vessel. A total of 320 VIM-IR cells were traced in this study, as four cells per region were reconstructed and analyzed for all three regions of all 20 subjects (where cortical grey and white matter were considered as independent compartments). Analyses were performed on all reconstructed cells with Neurolucida Explorer (MBF Bioscience) by an experimenter blinded to the group identity of each sample. Branched Structure Analysis (BSA) was used to compare seven structural features of astrocytes: process number, node number, terminal number, mean process length, total process length, mean process area, and soma area.

### 3.5 3D Reconstructions

All 3D reconstructions were created in Blender (Amsterdam, the Netherlands), which is a free, open-source 3D software suite. For the morphological reconstructions featured in Figure 3A, Neurolucida.DAT files were converted online into .SWC files for two morphological tracings of VIM-IR astrocytes in the prefrontal cortex white matter, which had process numbers representative of controls and depressed suicides in this region. These were then imported into Blender with the assistance of the Neuromorphovis plugin, and BSA features were manually colored and annotated for demonstration purposes (Abdellah *et al*., 2018). For the ‘stereological cube’ models featured in Figure 6, a randomly distributed particle system of 25mm^3^ spheres within the volume of 1m^3^ cube was used to visualize regional densities of 25µm^3^ diameter cell bodies counted within the regional brain volume of 1mm^3^. Sphere volume was kept constant across markers and regions to facilitate cell density comparisons.

### 3.6 Statistical analysis

All measurements are expressed as mean ± standard error of the mean (SEM), graphs display corrected p values on data after outlier removal, and p < 0.05 was considered significant in all statistical tests. Preliminary statistical analyses were performed using Prism v. 6.04 (GraphPad Software, San Diego, CA, USA). Data were assessed for a normal distribution using the Kolmogorov–Smirnov test. Statistical outliers were identified using a ROUT outlier test with Q = 1% (Motulsky & Brown, 2006). Three data points were identified as outliers in this study, one for VIM-IR astrocyte density in PFC GM of a depressed suicide, and two for GFAP-IR astrocyte density in the thalamus of depressed suicides. In these two instances we provide in-text the p value before outlier removal, as assessed by the Mann-Whitney test — outliers were removed to normalize both data sets, so they met parametric test assumptions. All other instances of uncorrected and corrected group differences were assessed for each region using unpaired t-tests using SPSS version 21 software (IBM Corporation), correction was made for three covariates — age, pH and PMI — to correct for any potential effects they may have had on density or morphometry measurements.

## 4 Results

### 4.1 GFAP-IR astrocyte densities are generally reduced in depressed suicides

Using a stereological approach to assess the regional densities of astrocytes in an unbiased fashion, GFAP-IR astrocytes were found to be lower in all regions for depressed suicides relative to controls (**Fig. 1A**). Prefrontal cortex grey matter had half as many GFAP-IR astrocytes in depressed suicides than in controls (444 ± 73 cells/mm^3^ vs. 1035 ± 232 cells/mm^3^; p [uncorrected] = 0.03, p [corrected] = 0.04). Prefrontal cortex white mater had almost half as many GFAP-IR astrocytes in depressed suicides than in controls, although this difference was not statistically significant (1353 ± 373 cells/mm^3^ vs. 2668 ± 510 cells/mm^3^; p [uncorrected] = 0.05, p [corrected] = 0.07). The mediodorsal thalamus had the greatest difference between groups, with depressed suicides displaying 5-fold fewer GFAP-IR astrocytes than depressed suicides (1049 ± 400 cells/mm^3^ vs. 5902 ± 1348 cells/mm^3^; p [uncorrected] = 0.003), but this became 12-fold after removing one statistical outlier that prevented parametric testing (491 ± 101 cells/mm^3^ vs. 5902 ± 1348 cells/mm^3^; p [uncorrected] = 0.003, p [corrected] = 0.005). Similarly, the caudate nucleus also had 5-fold fewer GFAP-IR astrocytes in depressed suicides than in controls (204 ± 87 cells/mm^3^ vs. 1142 ± 356 cells/mm^3^; p [uncorrected] = 0.02, p [corrected] = 0.03).

**Figure 1.**
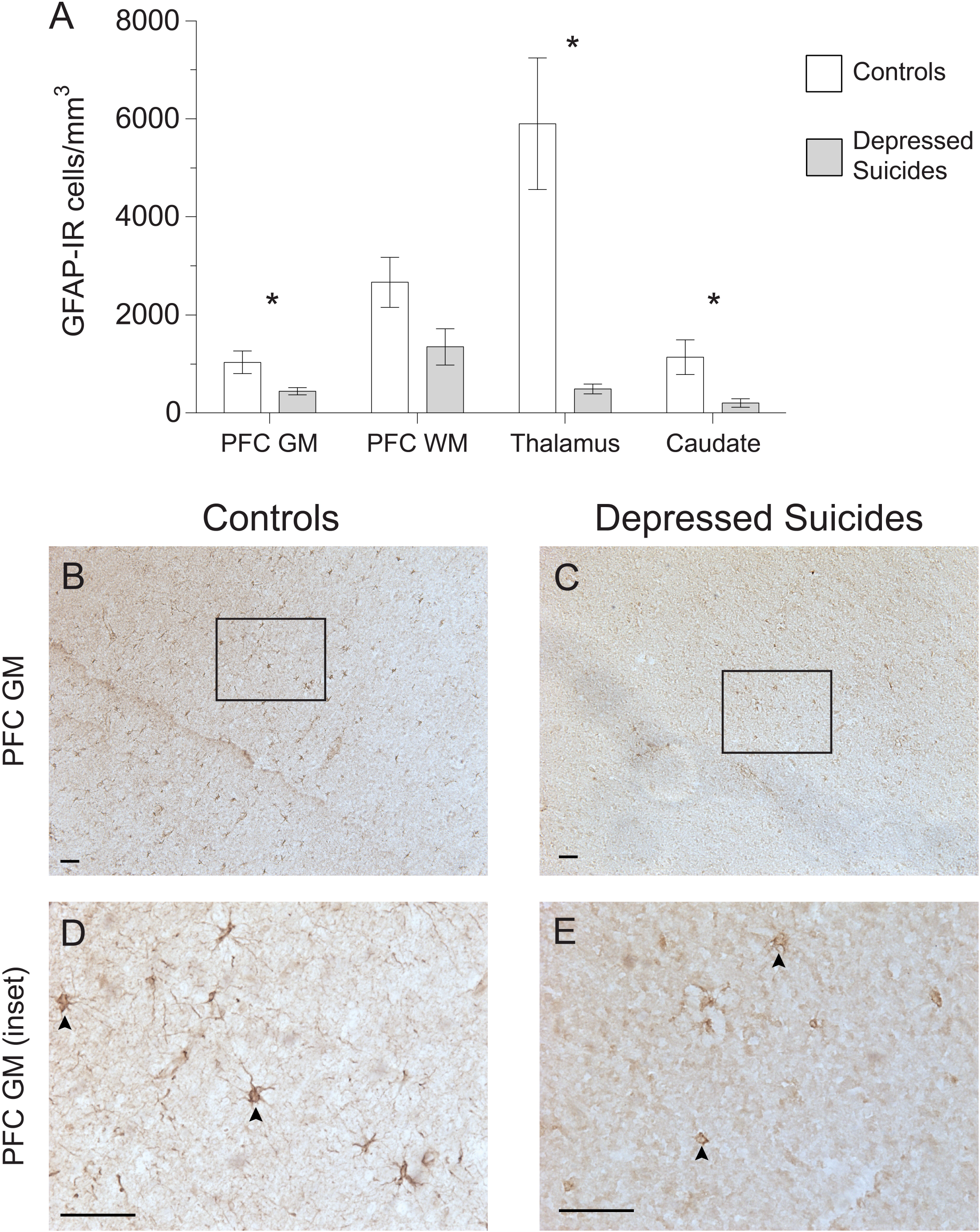
Lower densities of GFAP-IR cerebral astrocytes in depressed suicides relative to controls. **(A)** Representative micrographs illustrating GFAP-IR astrocytes in the prefrontal cortex gray matter (PFC GM). Scale bars = 50µm. **(B)** Depressed suicides had significantly lower densities of GFAP-IR astrocytes than controls in all regions examined except in the prefrontal cortex white matter (PFC WM), in which the difference was nearly significant. * p ≤ 0.05; n = 10; unpaired t-tests corrected for age, pH and postmortem interval.

### 4.2 Vimentin-IR astrocyte densities are generally reduced in depressed suicides

As the regional densities of GFAP-IR and VIM-IR were recently reported to differ in postmortem samples from healthy human brains (O’Leary *et al*., 2020), we next assessed whether VIM-IR astrocytes would also display altered regional densities in depressed suicides (**Fig. 2A**). Prefrontal cortex grey matter had 6-fold fewer VIM-IR astrocytes in depressed suicides than in controls (137 ± 100 vs. 886 ± 316 cells/mm^3^; p [uncorrected] = 0.08), and this difference became over 20-fold and statistically significant after removing one statistical outlier (38 ± 15 vs. 886 ± 316 cells/mm^3^; p [uncorrected] = 0.02, p [corrected] = 0.03). Prefrontal cortex white matter had 20-fold fewer VIM-IR astrocytes in depressed suicides than in controls (14 ± 5 vs. 278 ± 98 cells/mm^3^; p [uncorrected] = 0.002, p [corrected] = 0.001). There were slightly fewer VIM-IR astrocytes in the mediodorsal thalamus of controls relative to depressed suicides, but this was not statistically significant due to there being very few cells in both groups (1.2 ± 0.2 vs. 2.1 ± 0.5 cells/mm^3^; p [uncorrected] = 0.12, p [corrected] = 0.05). The caudate nucleus had 10-fold fewer VIM-IR astrocytes in depressed suicides than in controls (104 ± 38 vs. 1179 ± 355 cells/mm^3^; p [uncorrected] = 0.01, p [corrected] = 0.01).

**Figure 2.**
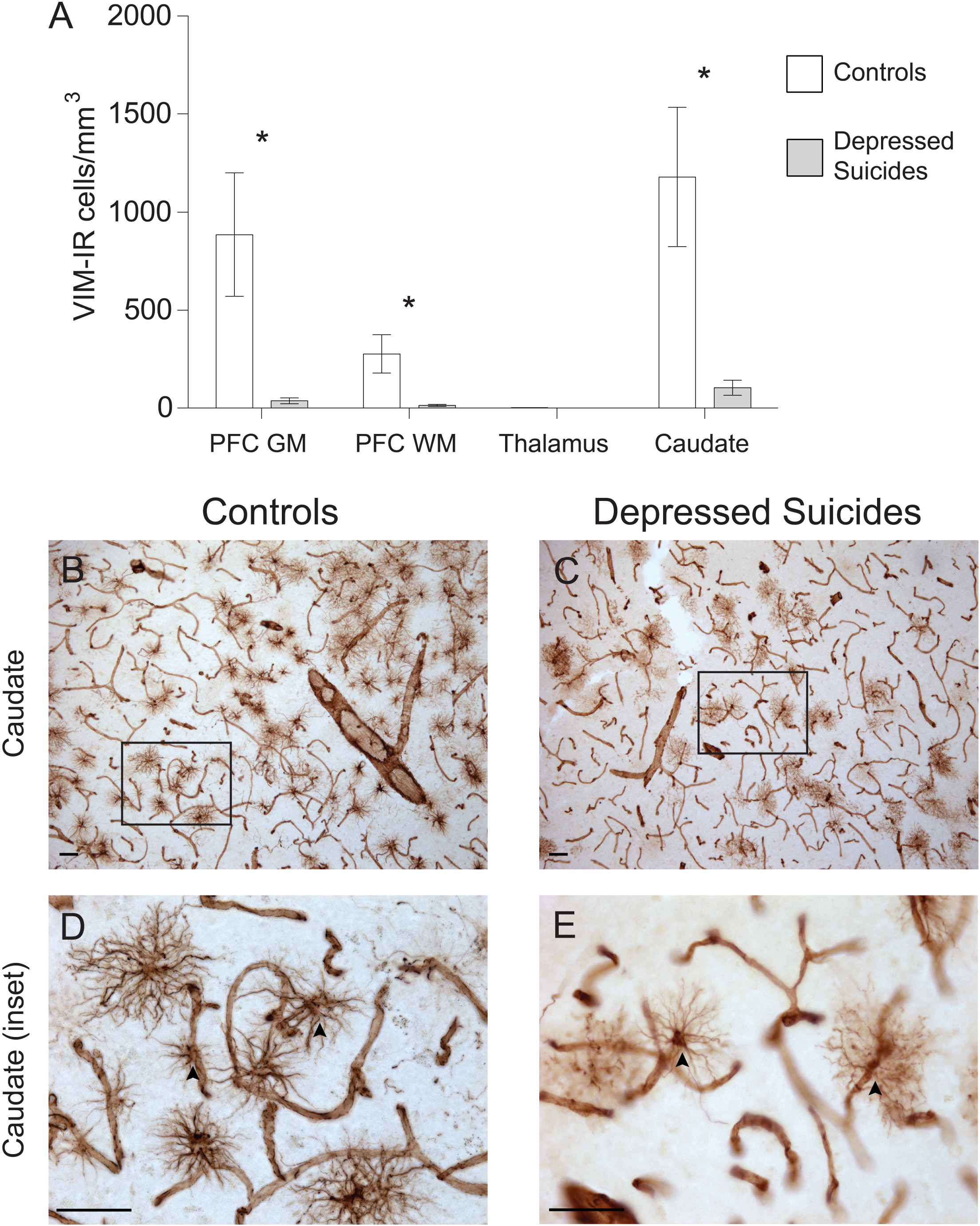
Lower densities of VIM-IR cerebral astrocytes in depressed suicides relative to controls. **(A)** Representative micrographs illustrating VIM-IR astrocytes in the caudate nucleus. Scale bars = 50µm. **(B)** Depressed suicides had significantly lower densities of GFAP-IR astrocytes than controls in the caudate nucleus, the prefrontal cortex gray matter (PFC GM) and the prefrontal cortex white matter (PFC WM). No group difference was observed in the mediodorsal thalamus, which presented exceedingly few VIM-IR astrocytes in both groups. * p ≤ 0.05; n = 10; unpaired t-tests corrected for age, pH and postmortem interval.

### 4.3 Vimentin-immunoreactive astrocytes: mostly unchanged morphology in depressed suicides

#### 4.3.1 Process structure

The morphology of VIM-IR astrocytes mostly did not differ significantly between depressed suicides and controls (**Fig. 3;** see parameters assessed in **3A**). The only statistically significant group difference for process number was in the prefrontal cortex white matter (**Fig. 3B**); relative to controls, astrocytes from depressed suicides had on average two more processes in prefrontal cortex grey matter (30 ± 1 vs. 28 ± 2 processes; p [uncorrected] = 0.60, p [corrected] = 0.83), six fewer processes in prefrontal white matter (29 ± 2 vs. 35 ± 1 processes; p [uncorrected] = 0.008, p [corrected] = 0.02), three fewer processes in the thalamus (18 ± 1 vs. 21 ± 1 processes; p [uncorrected] = 0.07, p [corrected] = 0.06), and the same number of processes in the caudate nucleus (27 ± 1 vs. 27 ± 1 processes; p [uncorrected] = 0.63, p [corrected] = 0.77). Astrocyte node number did not significantly differ between groups for any region (**Fig. 3C**); relative to controls, astrocytes from depressed suicides had on average three fewer nodes in prefrontal cortex grey matter (30 ± 3 vs. 33 ± 2 nodes; p [uncorrected] = 0.47, p [corrected] = 0.26), two fewer nodes in prefrontal white matter (20 ± 2 vs. 22 ± 2 nodes; p [uncorrected] = 0.50, p [corrected] = 0.59), the same number of nodes in the thalamus (14 ± 1 vs. 14 ± 2 nodes; p [uncorrected] = 0.85, p [corrected] = 0.95), and one less node in the caudate nucleus (32 ± 3 vs. 33 ± 3 nodes; p [uncorrected] = 0.82, p [corrected] = 0.91). Similarly, terminal number was not significantly affected in depressed suicides (**Fig. 3D**); relative to controls, astrocytes from depressed suicides had on average two fewer terminals in prefrontal cortex grey matter (60 ± 4 vs. 62 ± 4 terminals; p [uncorrected] = 0.70, p [corrected] = 0.43), eight fewer terminals in prefrontal white matter (49 ± 2 vs. 57 ± 3 terminals; p [uncorrected] = 0.05, p [corrected] = 0.08), two fewer terminals in the thalamus (33 ± 2 vs. 35 ± 2 terminals; p [uncorrected] = 0.39, p [corrected] = 0.33), and the same number of terminals in the caudate nucleus (61 ± 3 vs. 61 ± 3 terminals; p [uncorrected] = 0.94, p [corrected] = 0.87).

**Figure 3.**
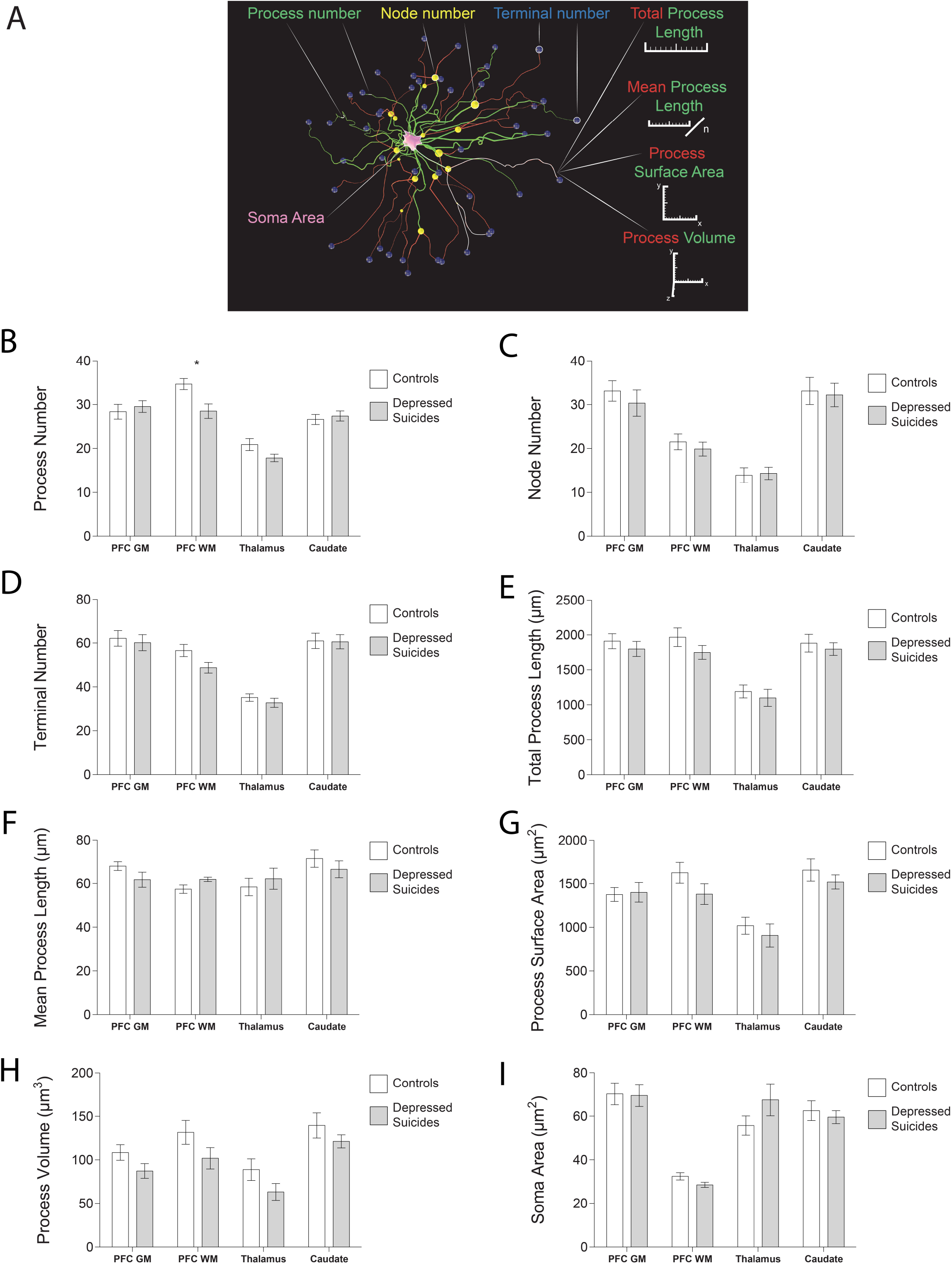
VIM-IR astrocyte morphology is generally similar in depressed suicides vs controls. (**A)** 3D reconstruction of a VIM-IR astrocyte from the prefrontal cortex white matter (PFC WM) representative of those from depressed suicides. The soma (pink) that extends primary processes (green) which can branch at nodes (yellow) into secondary processes (red) that eventually end as terminals (blue). Branched Structure Analysis (BSA) measurements have been annotated. **(B–H)** The BSA revealed a lower (primary) process number for VIM-IR astrocytes in the PFC WM of depressed suicides relative to controls. There were no group differences for any BSA measurements of VIM-IR astrocytes in the prefrontal cortex gray matter (PFC GM), thalamus or caudate nucleus. *p ≤ 0.05; n = 10; unpaired t-tests corrected for age, pH and postmortem interval.

#### 4.3.2 Process length

Astrocyte total process length did not significantly differ between groups in any region (**Fig. 3E**); relative to controls, the mean total process length for astrocytes from depressed suicides was consistently lower in the prefrontal cortex grey matter (1800 ± 108 vs. 1911 ± 108 µm; p [corrected] = 0.47, p [corrected] = 0.29), the prefrontal white matter (1750 ± 98 vs. 1968 ± 133 µm; p [uncorrected] = 0.21, p [corrected] = 0.29), the thalamus (1100 ± 121 vs. 1192 ± 92 µm; p [uncorrected] = 0.55, p [corrected] = 0.46), and the caudate nucleus (1798 ± 91 vs. 1882 ± 126 µm; p [uncorrected] = 0.59, p [corrected] = 0.79). Across all regions astrocyte mean process length did not significantly or consistently differ between groups (**Fig. 3F**); relative to healthy controls, astrocyte processes from depressed suicides were shorter in prefrontal cortex grey matter (62 ± 3 vs. 68 ± 2 µm; p [uncorrected] = 0.14, p [corrected] = 0.14), longer in prefrontal white matter (62 ± 1 vs. 57 ± 2 µm; p [uncorrected] = 0.06, p [corrected] = 0.08), longer in the thalamus (62 ± 5 vs. 58 ± 4 µm; p [uncorrected] = 0.56, p [corrected] = 0.73) and shorter in the caudate nucleus (67 ± 4 vs. 71 ± 4 µm; p [uncorrected] = 0.39, p [corrected] = 0.53).

#### 4.3.3 Cell size

The surface area of astrocyte processes did not significantly differ between groups for any region (**Fig. 3G**); relative to controls, the surface area of astrocyte processes from depressed suicides was smaller in the prefrontal cortex grey matter (1377 ± 79 vs. 1401 ± 112 µm^2^; p [uncorrected] = 0.86, p [corrected] = 0.80), the prefrontal white matter (1381 ± 119 vs. 1625 ± 119 µm^2^; p [uncorrected] = 0.16, p [corrected] = 0.20), the thalamus (907 ± 133 vs. 1019 ± 98 µm^2^; p = [uncorrected] 0.50, p [corrected] = 0.47), and the caudate nucleus (1519 ± 81 vs. 1657 ± 128 µm^2^; p [uncorrected] = 0.38, p [corrected] = 0.53). As for total process length, the volume occupied by astrocyte processes did not significantly differ between group in any region (**Fig. 3H**), but relative to healthy controls, the mean volume of astrocyte processes from depressed suicides was consistently smaller in the prefrontal cortex grey matter (87 ± 8 vs. 108 ± 9 µm^3^; p [uncorrected] = 0.25, p [corrected] = 0.24), the prefrontal white matter (102 ± 12 vs. 132 ± 14 µm^3^; p [uncorrected] = 0.30, p [corrected] = 0.16), the thalamus (63 ± 10 vs. 89 ± 12 µm^3^; p [uncorrected] = 0.31, p [corrected] = 0.30), and the caudate nucleus (121 ± 8 vs. 140 ± 14 µm^3^; p [uncorrected] = 0.32, p [corrected] = 0.25). The soma area of astrocytes did not significantly differ between groups for any region (**Fig. 3I**); relative to healthy controls, the soma area of astrocytes from depressed suicides was smaller in the prefrontal cortex grey matter (69 ± 5 vs. 70 ± 5 µm^2^; p [uncorrected] = 0.92, p [corrected] = 0.76), smaller in the prefrontal white matter (29 ± 1 vs. 32 ± 2 µm^2^; p [uncorrected] = 0.07, p [corrected] = 0.06), larger in the thalamus (68 ± 7 vs. 56 ± 4 µm^2^; p [uncorrected] = 0.19, p [corrected] = 0.24), and smaller in the caudate nucleus (60 ± 3 vs. 62 ± 5 µm^2^; p [uncorrected] = 0.60, p [corrected] = 0.50).

### 4.4 CD31-immunoreactive vascular density is increased in the prefrontal cortex white matter of depressed suicides

There was no general trend for group differences in vascular density (**Fig. 4A**). In prefrontal cortex grey matter, it was found to be comparable between depressed suicides and controls (6.9% ± 0.6 vs. 7.4% ± 0.5% coverage; p [uncorrected] = 0.55, p [corrected] = 0.76). Similar results were found in the mediodorsal thalamus and caudate nucleus, in which vascular densities were also comparable between depressed suicides and controls (thalamus: 7.2% ± 0.4% vs. 7.5% ± 0.4% coverage; p [uncorrected] = 0.58, p [corrected] = 0.96; caudate: 8.1% ± 0.3% vs. 7.7% ± 0.5% coverage; p [uncorrected] = 0.46, p [corrected] = 0.28). Only in prefrontal cortex white matter was there a significant group difference in vascular density, with greater vascularization in depressed suicides than in controls (3.4% ± 1.2% vs. 2.6% ± 0.2% coverage; p [uncorrected] = 0.003, p [corrected] = 0.002).

**Figure 4.**
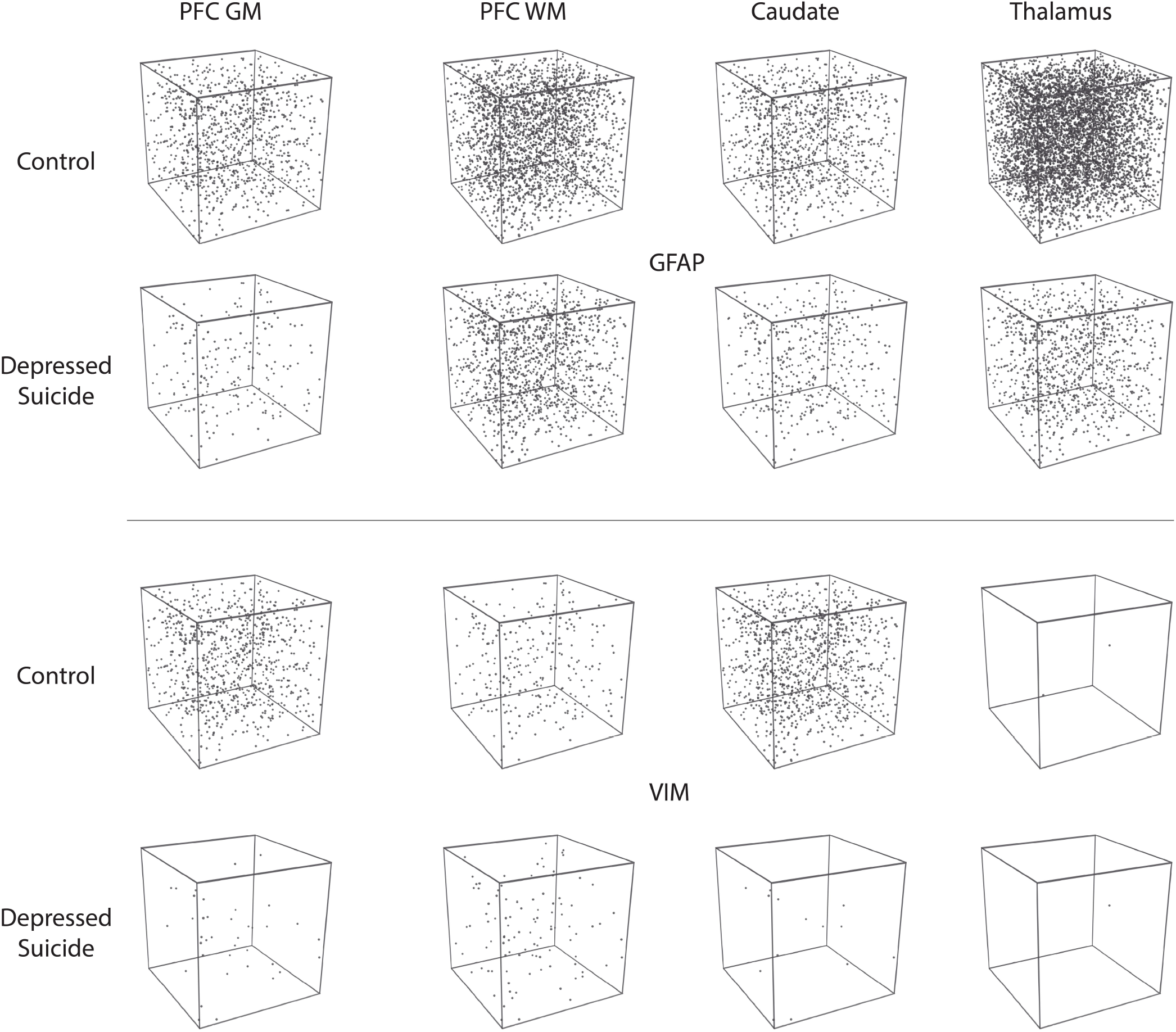
Increased CD31-IR vascular density in cortical white matter from depressed suicides relative to controls. **(A)** Representative micrographs showing that CD31-IR vascular density in the prefrontal cortex white matter (PFC WM). Scale bars = 50µm. **(B)** Depressed suicides had a significantly higher vascular density than controls in the mediodorsal thalamus. No group differences were observed in the prefrontal cortex gray matter (PFC GM), the caudate nucleus or the mediodorsal thalamus. * p ≤ 0.05; n = 10; unpaired t-tests corrected for age, pH and postmortem interval.

## 5 Discussion

To our knowledge, this is the first study investigating VIM-IR astrocytes in MDD and suicide, and the first cross-regional study of astrocyte density or morphology in MDD. This is one of few postmortem studies comparing the same samples with more than one astrocytic markers. The main results (1) further support previous reports of greater differences in GFAP expression in subcortical regions than in cortical regions of depressed suicides (Torres-Platas et al., 2016); (2) show that VIM-IR astrocyte densities are even more robustly decreased (without morphological changes) in cortical regions; (3) reveal altered CD31-IR vascular density in the PFC WM of depressed suicides. Presenting our data as densities using a stereological approach avoided the potential bias in counting from cortical volumes altered by depression or other factors. As depression generally decreases cortical volumes (Grieve et al., 2013), a non-stereological approach would likely give rise to increased or unchanged cortical astrocyte densities in case samples. However, previous stereological postmortem studies have reported decreased glial densities in the prefrontal cortex in depression (Rajkowska et al., 1999; Cotter et al., 2002), where we now report a decreased density of VIM-IR astrocytes.

### 5.1 Astrocyte densities

In this study, we revealed lower GFAP-IR and VIM-IR astrocyte densities in both cortical and subcortical brain samples from well-characterized depressed suicides compared to matched controls **(Fig. 5)**. For GFAP-IR astrocytes, this decrease was statistically significant in all regions except for the prefrontal cortex white matter (p = 0.053). This reveals a widespread alteration in GFAP-IR astrocytes throughout a number of cerebral networks in depressed individuals. The density of VIM-IR astrocytes was also assessed in the same subjects and regions. As our previous study showed that VIM mostly labels a subset of GFAP-IR astrocytes (O’Leary *et al*., 2020), this approach helped clarify whether reduced GFAP-IR astrocyte densities in depressed suicides reflects a reduction in astrocyte density as opposed to a reduction of GFAP immunoreactivity. In depressed suicides, the density of VIM-IR astrocytes was strongly and significantly lower in the prefrontal cortex and caudate nucleus, but not in the mediodorsal thalamus, in which exceedingly rare VIM-IR astrocytes are observed in controls. The lower densities of both GFAP-IR and VIM-IR astrocytes favors a hypothesis of a reduced number of astrocytes over a reduced reactive profile of astrocytes in depressed suicides.

**Figure 5.**
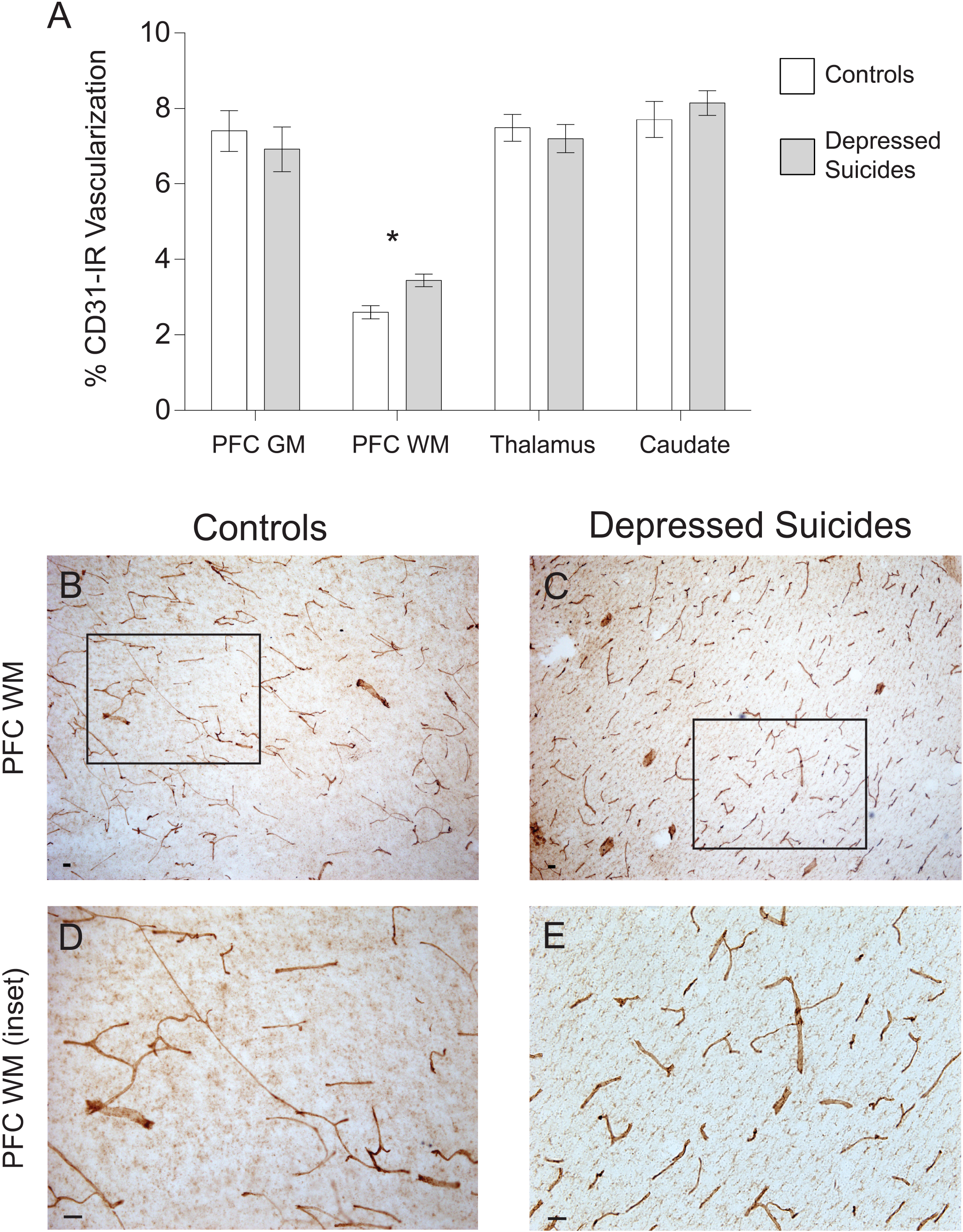
Visual representation of astrocyte densities in depressed suicides. Each cube represents 1mm^3^ of cerebral tissue in which are distributed the stereological estimates of cell densities reported in Figures 1 and 2 (cell body diameter = 25µm). These illustrations demonstrate that astrocyte density is greatly and widely affected in depressed suicides relative to controls.

The brain regions examined in this study are closely associated in a prominent frontal-subcortical circuit implicated in executive function — neurons project from the caudate nucleus to the mediodorsal thalamus and then to the dorsomedial prefrontal cortex gray matter (Bonelli & Cummings, 2007). This executive circuit has been found to be dysregulated in depression, with patients displaying a relatively lower resting state connectivity between the prefrontal cortex and either the striatum or the thalamus are more likely to have a positive treatment response to repetitive transcranial magnetic stimulation of the prefrontal cortex (Salomons *et al*., 2014). The connectivity of the prefrontal cortex seems essential to the neuroanatomy of depression — no single brain region consistently associates with MDD when lesioned, however, brain regions with relatively high resting state connectivity to the dorsomedial prefrontal cortex are consistently associated with MDD following lesions (Padmanabhan *et al*., 2019). In the context of our results, astrocyte dysfunction may directly regulate resting state functional connectivity to the prefrontal cortex, as astrocytes mediate vasodilation responsible for changes in regional cerebral blood flow. If our results do represent a reduction in numbers of astrocytes, this may represent a compensatory mechanism by which the brain is indirectly reducing functional connectivity by impairing the efficiency of gliovascular coupling in a circuit that becomes hyperconnected in depression.

A reduction in GFAP-IR astrocyte density in multiple brain regions from depressed suicides supports a strong consensus in the literature for decreased regional GFAP protein levels in postmortem studies and animal models of depression (Kim *et al*., 2018). The difference in GFAP-IR astrocyte density was greater in subcortical regions than in the cortical regions, which is in agreement with a previous report of GFAP protein levels being even more downregulated in the mediodorsal thalamus and caudate nucleus than in cortical regions of depressed suicides relative to controls (Torres-Platas *et al*., 2016). This previous study also found the mediodorsal thalamus and caudate nucleus have greater regional GFAP expression levels than many cortical regions in healthy adults, and so our findings indicate that brain regions that normally have a high GFAP expression in healthy adults may have particularly strong depression-related changes in GFAP-IR astrocyte densities. The association between regional GFAP expression and GFAP-IR astrocyte density is supported by a previous study that correlated low regional GFAP protein levels with low GFAP-IR area fraction in the dorsomedial prefrontal cortex gray matter of depressed suicides (Si *et al*., 2004), which may well resemble a reduction in cell density given our findings. Although we found no significant difference in GFAP-IR astrocyte density in dorsomedial prefrontal cortex white matter, a previous study reported a significantly lower density of GFAP-IR astrocytes in the ventromedial prefrontal cortex white matter of depressed suicides — a difference that was 5-fold greater than was observed for dorsomedial prefrontal cortex white matter in the present study (Rajkowska *et al*., 2018). This suggests that prefrontal cortex white matter may have a particularly strong subregional heterogeneity for depression-related changes in GFAP-IR astrocyte density.

While VIM-IR astrocytes clearly represent a minority of the total astrocyte population in healthy controls (O’Leary et al., 2020), VIM is a valuable marker as it is strongly immunoreactive in both cell body and processes. The group differences in astrocyte density in the prefrontal cortex grey matter and caudate nucleus were twice as great for VIM-IR astrocytes than for GFAP-IR astrocytes, indicating astrocytes strongly expressing VIM may be especially affected in depressed suicides. The lack of a significant decrease of VIM-IR astrocyte density in the mediodorsal thalamus was unsurprising, given that many sections from the thalamus of control subjects in this and our previous study contained no VIM-IR astrocytes at all. However, we suspect the consistent paucity of VIM-IR astrocytes in the mediodorsal thalamus in health and illness indicates an unrecognized functional role of astrocytes in the thalamus that does not require VIM expression. This is further suggested by the substantial differences in astrocyte density in different nuclei of the thalamus in mouse brain (Emsley & Macklis, 2006).

To date, only one other postmortem study has assessed VIM in the context of depression and found no significant decrease of VIM mRNA levels in MDD in the anterior cingulate cortex of depressed individuals who died by natural causes (not by suicide) (Qi *et al*., 2019). Although this may reflect an effect unique to the anterior cingulate cortex, this does not reliably imply no change of VIM-IR astrocyte density in the ACC, given that the most VIM expression in the brain is found in vascular endothelial cells (Nagy *et al*., 2020; O’Leary et al., 2020). As for astrocytes labelled with glutamine synthetase, there is a reduced number of VIM-IR astrocytes in postmortem brain samples from individuals with MDD, but due to its expression in other cell types, this effect is not detected at the regional level of protein or mRNA (Bernstein *et al*., 2015). The reduction of VIM-IR astrocyte density in depressed suicides suggests that reduced GFAP levels do not correspond specifically to GFAP dysfunction, but rather a widespread loss or dysfunction of astrocytes. A pan-astrocytic marker like Aldh1L1 will be needed to infer whether only reactive astrocytes expressing VIM and GFAP are affected in depression; however, this seems unlikely, as genes specific to both non-reactive and reactive astrocytes are downregulated in many brain regions in depressed suicides (Nagy *et al*., 2015; Nagy *et al*., 2017; Zhao *et al*., 2016) and non-depressed suicides (Zhao *et al*., 2018).

### 5.2 Vascular densities

In a previous study, our group showed that regional astrocyte density inversely correlates with regional vascular density in postmortem brain samples from controls (O’Leary *et al*., 2020). In the present study, brain regions from depressed suicides had significantly lower astrocyte density than controls, and so we anticipated that they may also have a correspondingly greater vascular density. Depressed suicides had a significantly different (higher) vascular density than controls only in prefrontal cortex white matter— no other significant differences or general trends were observed for regional vascular density. This finding suggest that gliovascular interactions in cortical white matter may be preferentially affected by depression, as also suggested by a previous report of reduced coverage of blood vessels by astrocytic endfeet in prefrontal cortex grey matter, but not white matter (Rajkowska *et al*., 2013). It is also likely that vascular density does not reflect the full extent of vascular changes in depressed individuals, such as the known reduction in claudin-5 protein levels in postmortem vascular endothelial cells of the nucleus accumbens in depressed suicides, which in animal models has been associated with an abnormally increased permeability of the blood brain barrier (Menard *et al*., 2017; Dudek *et al*., 2020). Future studies on gliovascular interactions in depression may need to resolve this with finer precision with a non-stereological method, given the absence of a clear region-wide relationship between astrocyte and vascular density in depression.

### 5.3 Limitations and Future Directions

The two main limitations of this study were its relatively small sample size and lack of samples from females. Female samples were not included due to the low availability of such samples from depressed suicides which could be effectively matched for factors that are known to greatly affect astrocyte regional densities and gene expression in postmortem samples, including age and postmortem interval (Miguel-Hidalgo *et al*., 2000; Nagy *et al*., 2015).

In conclusion, different brain regions in depressed suicides exhibit robust reductions in VIM-IR astrocyte densities that are even greater than those for GFAP-IR astrocytes. Our data also more generally revealed a consistent cross-regional trend for reduced astrocyte densities in depressed suicides, and a unique change in vascular density in the prefrontal cortex white matter. With the minor exception of fibrous astrocytes in the prefrontal cortex, there were no clear changes in the morphology of VIM-IR astrocytes that we could most clearly and precisely observe, indicating that depression has a larger and more widespread effect on astrocyte density than on astrocyte morphology in mood-associated brain regions. In the future, single cell sequencing approaches will be informative for establishing whether the function of these smaller populations of cells in depressed suicides are altered in depressed suicides.

## 7 Acknowledgments

The present study used the services of the Douglas-Bell Canada Brain Bank and of the Molecular and Cellular Microscopy Platform at the Douglas Hospital Research Centre. The authors are grateful to Maâmar Bouchouka, Josée Prud’homme, Dominique Mirault and Melina Jaramillo Garcia for their kind assistance.

## 8 Author Contributions

LAO and NM conceived the project, designed the experiments and drafted the manuscript. CB, MAD and LAO conducted tissue processing, and LAO conducted the IHC experiments. LAO and JCM performed the cell counting experiments. LAO performed vascular density and astrocyte morphometry experiments and analyzed all the data. All authors contributed to the interpretation of the results in addition to participating in the finalization of the manuscript.

## 10 Contribution to the Field Statement

Astrocytes are a glial cell type in the brain that are less abundant in the brains of depressed individuals. It is unclear exactly which astrocyte populations or brain regions are most affected in depression, because most previous studies have studied only one astrocyte population in one brain region in the same individuals. It is also unclear whether the reduced number of astrocytes during depression is accompanied with changes to their physical structure or to the blood vessels they normally contact. This study is also the first anatomical comparison of astrocytes labelled by the protein vimentin in postmortem brain tissue from depressed individuals. We performed the first cross-regional assessment of astrocyte density and morphometry using two distinct astrocytic markers, as well as of vascular density in postmortem brain samples from depressed individuals and matched controls. We found a significant and widespread reduction in astrocyte densities (but no significant change in cell morphology) in depressed suicides relative to controls. Blood vessel density was only different (increased) in prefrontal cortex white matter of depressed suicides. This study reveals a widespread alteration of cerebral astrocytes in depression and suicide.

